# A Developmental Reduction of the Excitation:Inhibition Ratio in Association Cortex during Adolescence

**DOI:** 10.1101/2021.05.19.444703

**Authors:** Bart Larsen, Zaixu Cui, Azeez Adebimpe, Adam Pines, Aaron Alexander-Bloch, Max Bertolero, Monica E. Calkins, Raquel E. Gur, Ruben C. Gur, Arun S. Mahadevan, Tyler M. Moore, David R. Roalf, Jakob Seidlitz, Valerie J. Sydnor, Daniel H. Wolf, Theodore D. Satterthwaite

**Author notes:** Authors contributed equally (Daniel H. Wolf, Theodore D. Satterthwaite).

## Abstract

Adolescence is hypothesized to be a critical period for the development of association cortex. A reduction of the excitation:inhibition (E:I) ratio is a hallmark of critical period development; however it has been unclear how to assess the development of the E:I ratio using non-invasive neuroimaging techniques. Here, we used pharmacological fMRI with a GABAergic benzodiazepine challenge to empirically generate a model of E:I ratio based on multivariate patterns of functional connectivity. In an independent sample of 879 youth (ages 8-22 years), this model predicted reductions in the E:I ratio during adolescence, which were specific to association cortex and related to psychopathology. These findings support hypothesized shifts in E:I balance of association cortices during a neurodevelopmental critical period in adolescence.

**Teaser:** Inhibitory maturation of the association cortex reflects an adolescent critical period.

## Introduction

Adolescent brain development is characterized, in part, by the continued structural and functional maturation of the association cortices (*1*–*8*). The specificity of the developmental timing and localization of association cortex maturation as well as the links between association cortex development and long-term psychiatric outcomes have led to the hypothesis that adolescence functions as a critical period of development within association cortex (*9*, *10*). Critical periods are windows of development during which experience powerfully shapes the development of neural circuits through heightened experience-dependent plasticity with long-term impacts on behavior (*11*). These important neurodevelopmental windows are theorized progress hierarchically throughout development, beginning in primary sensory cortices and sequentially advance to secondary and higher-order cortical areas (*11*, *12*). The neurobiological mechanisms that underlie critical periods are thought to be conserved across the cortex and have been carefully delineated in decades of work on early critical periods in sensory cortex (*11*–*14*).

One of the hallmark features of critical period development is the maturation of GABAergic inhibitory circuitry, particularly parvalbumin positive interneurons, leading to a reduction in the excitation to inhibition (E:I) ratio (*13*, *15*). The reduction of the E:I ratio leads to an increase in the signal-to-noise ratio of local circuit activity as inhibition suppresses the effect of spontaneous activity on circuit responses in favor of stimulus-evoked activity (*16*). This essential mechanism has been shown to regulate the timing of critical period development across visual (*14*), auditory (*17*), and sensorimotor cortices (*18*). As such, if the adolescent critical period hypothesis is correct, inhibitory maturation should result in a developmental reduction in the E:I ratio across adolescence within association cortex.

Evidence for E:I development in association cortex during adolescence has been largely limited to animal models. This work has suggested prefrontal GABAergic inhibitory circuitry undergoes significant modifications. Specifically, parvalbumin (PV) interneurons, a critical component E:I maturation in sensory system critical periods, have been shown increase in prefrontal cortex during adolescence in the rat (*19*) and non-human primate (*20*, *21*). At the same time, the expression of GABA_A_ receptor α1 subunits, which are primarily expressed on PV cells and support fast synaptic inhibition (*22*) as well as synaptic plasticity (*23*), also increase during adolescence in the prefrontal cortex of the non-human primate (*24*, *25*). These neurobiological changes lead to important functional increases in inhibitory signaling, effectively reducing the E:I ratio(*26*, *27*). Together, these findings are suggestive of critical period development and may indicate similar processes are unfolding in the human (*28*). Translating these findings to human studies of development is crucial as disruptions to the E:I balance are hypothesized to play a significant role in the onset of psychiatric disorders (*29*–*31*). However, the extent to which these critical period mechanisms are present in association cortex during adolescence in the human remains largely unexplored. Corroborating evidence has been found in postmortem studies which demonstrate increases in PV (*32*) and GABA_A_ α1 expression (*33*), but it has been unclear how to measure developmental changes in the E:I ratio *in vivo* in humans using available neuroimaging techniques. This lack of *in vivo* measures has limited our ability to test the adolescent critical period hypothesis.

Here, we leveraged a pharmacological fMRI (phMRI) experiment using a GABAergic benzodiazepine challenge to empirically generate a model for the effect of inhibitory modulation of the E:I ratio on patterns of fMRI connectivity. Benzodiazepines are positive allosteric modulators of the GABA_A_ receptor that increase the effectiveness of post-synaptic GABAergic signaling, resulting in an increase in inhibition relative to excitation. Benzodiazepines have been used to pharmacologically manipulate the E:I ratio by enhancing inhibitory signaling in disease models of E:I imbalance (*34*–*36*) as well as in studies of critical period development (*13*, *23*, *37*). In the current study, we first trained a multivariate model to distinguish benzodiazepine-induced change in the E:I ratio and established the neurobiological relevance of our empirical model by comparing the model features to known aspects of benzodiazepine pharmacology as well as a functional gradient that has been shown to reflect patterns of excitatory and inhibitory interneuron expression (*38*, *39*). We then applied our trained and validated model to a large, independent developmental dataset to investigate E:I changes occurring in association cortex during adolescence. We hypothesized that patterns of functional connectivity would develop to reflect a reduction in the E:I ratio that is specific to association cortex.

## Results

### An empirical model of the E:I ratio

Forty-three adult participants completed a double-blind, placebo-controlled phMRI study with the benzodiazepine alprazolam (86 sessions total). Alprazolam is a classical benzodiazepine that enhances the effect of GABA at GABA_A_ receptors through positive allosteric modulation, increasing inhibition and effectively reducing the E:I ratio (*40*). Functional connectivity matrices were derived for placebo and drug phMRI sessions using a top performing pipeline that minimized the impact of motion artifact (*41*). A linear support vector machine (SVM) classifier was trained to distinguish placebo and drug sessions based on the multivariate patterns of functional connectivity (**Figure 1**, green pathway). Cross-validation and permutation testing revealed that the trained SVM identified drug vs. placebo sessions in left-out data far better than chance (AUC = .716, *p_permutation_* = .002; **Figure 2a**). Sensitivity analyses confirmed that in-scanner head motion was not associated with our pharmacological manipulation or model performance (**Supplemental Figure 1**). The spatial pattern of estimated feature weights from the SVM model highlighted the contributions of subcortical regions, including the thalamus and amygdala, and also contributions throughout the cortex (**Figure 2b**).

**Figure 1.**
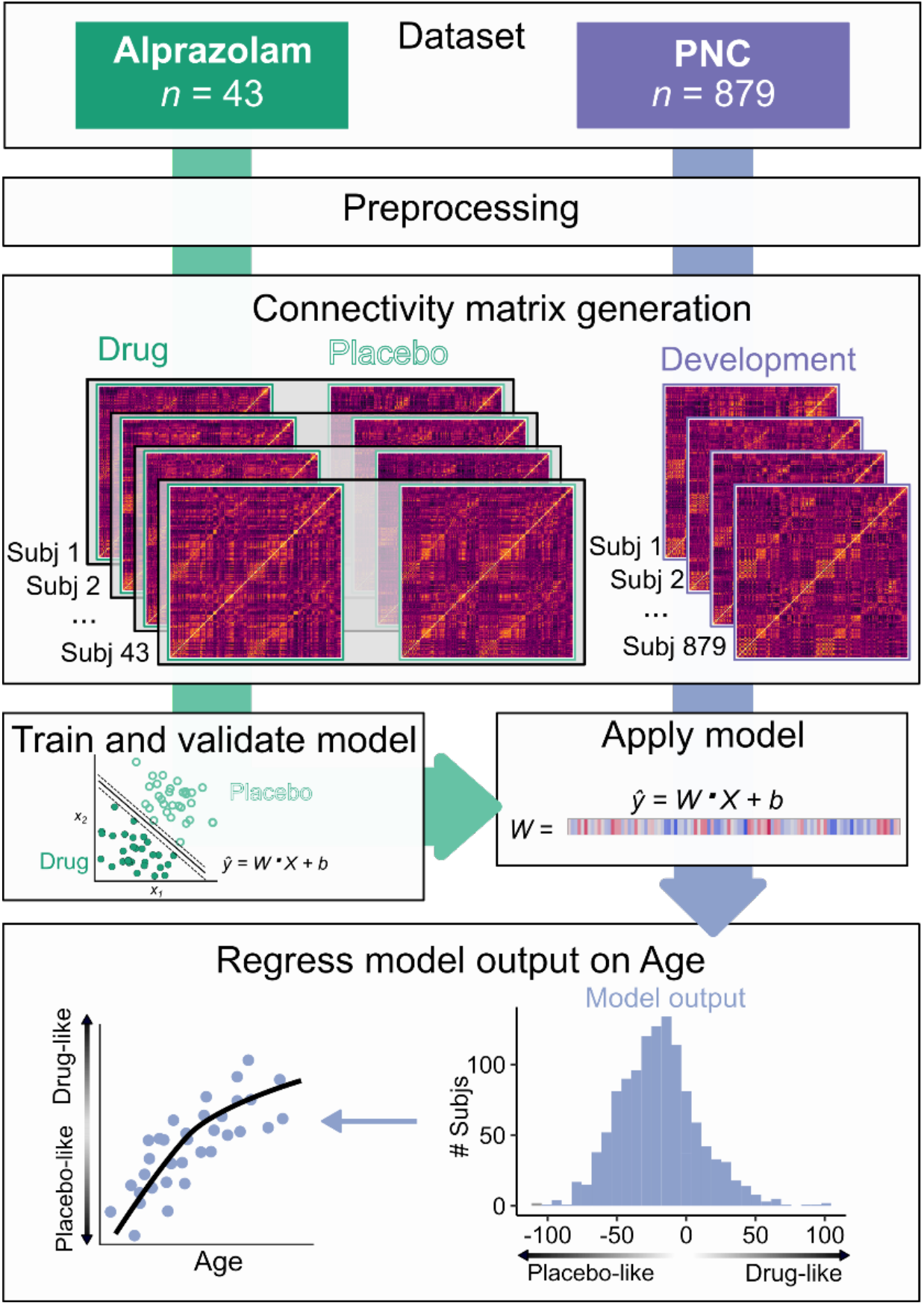
Analysis workflow. *Dataset:* Two datasets were collected on the same scanner using highly similar acquisition parameters: a phMRI dataset using the benzodiazepine alprazolam (green) and a developmental fMRI sample from the Philadelphia Neurodevelopmental Cohort (PNC; purple). *Preprocessing:* Datasets were preprocessed using identical pipelines which included removal of nuisance signal with aCompCor (, global signal regression, and task regression. *Connectivity matrix generation:* Connectivity matrices were generated from standard atlases for placebo and drug sessions from the alprazolam dataset (*n* = 43; 86 sessions total) and for the PNC dataset (*n* = 879). *Train and validate model:* The alprazolam dataset was used to train a linear SVM classifier to distinguish drug and placebo sessions using 10-fold cross-validation. *Apply model:* The validated alprazolam model was applied to the PNC dataset, generating a distance metric that reflected each participant’s position on a continuum from “drug-like” (lower E:I) to “placebo-like” (higher E:I). *Regress model output on age:* This metric was then regressed on age using a generalized additive model with penalized splines that included covariates for sex, head motion, and attentiveness.

**Figure 2.**
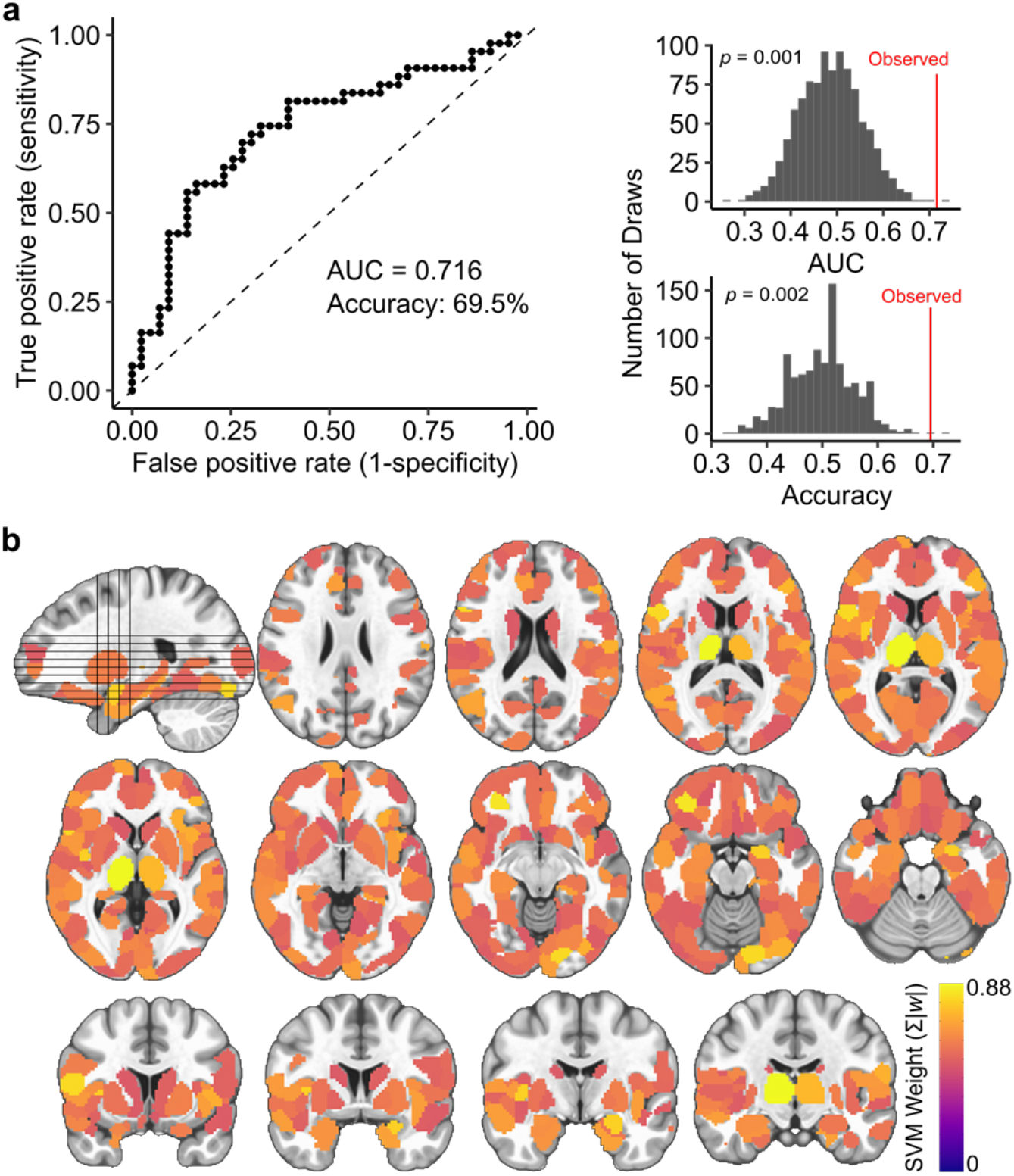
A multivariate model distinguishes alprazolam and placebo sessions, capturing E:I ratio. **a)** Classifier performance. The binary SVM classifier identified drug and placebo sessions in 10-fold cross-validation with an AUC of .716 and an accuracy = 69.5% (top). The observed AUC and accuracy were significantly greater than a permuted null distribution (bottom). **b)** Mean absolute feature weights for all nodes from the validated SVM model.

### Biological relevance of the E:I model

Next, we established the biological relevance of the trained E:I model. First, we compared the spatial pattern of cortical feature weights to a widely used functional gradient of macroscale cortical organization that places regions on a continuum from unimodal to transmodal function (*42*). This continuum has been shown to capture variation in excitatory neuron structure, inhibitory interneuron expression, and E:I balance (*38*, *39*). Using a recently-developed analytic procedure that accounts for spatial autocorrelation structure (*43*), we observed a significant relationship between our model feature weights and this pattern of macroscale cortical organization (*r* = .33; *p* = .003; **Figure 3a**). This finding suggests that GABAergic modulation of functional connectivity patterns varies along a transmodal-to-unimodal gradient that in part indexes diversity in excitatory and inhibitory neurobiological properties. Next, we investigated whether the estimated model features corresponded to the known pharmacology of benzodiazepines like alprazolam. Of the six GABA_A_ subunit receptors, GABA_A_ α1–6, only α1, α2, α3, and α5 are sensitive to benzodiazepines due to the presence of an amino acid residue, histidine (*44*, *45*). We used the Allen Human Brain Atlas (*46*) to evaluate how the feature weights from the classifier model aligned with spatial patterns of gene expression for the six GABA_A_ subunit receptors, GABR_A_1–6 (corresponding to GABA_A_ α1-6). We found evidence of a clear biological double dissociation: model features were significantly associated with the gene expression patterns of the benzodiazepine-sensitive GABA_A_ subunits (α1,α2,α3,α5) and <u>not</u> the benzodiazepine-*in*sensitive GABA_A_ subunits (α4, α6; **Figure 3b**)(*47*, *48*).

**Figure 3.**
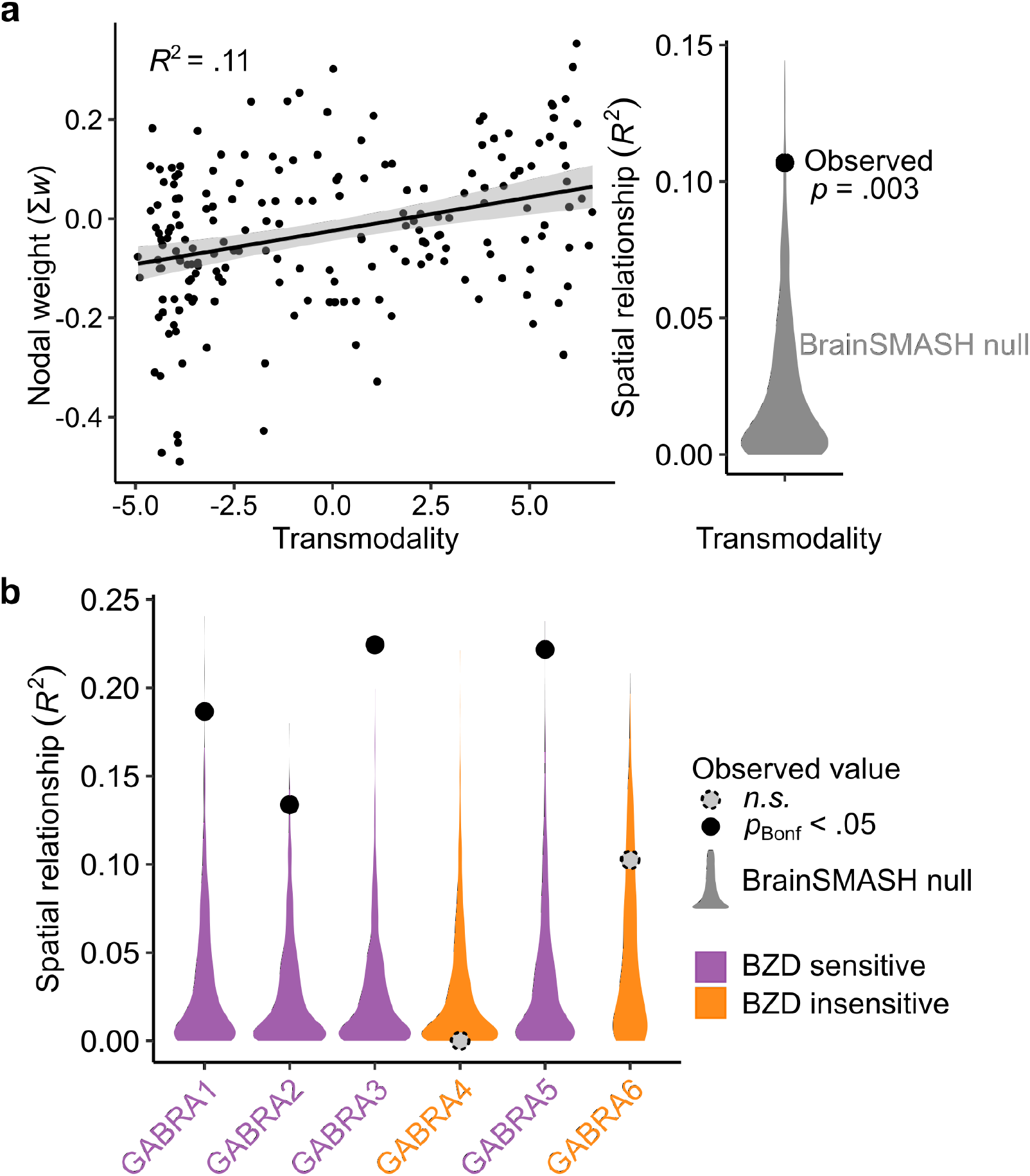
Model features align with cortical organization and benzodiazepine pharmacology. **a)** The cortical pattern of nodal SVM weights from the multivariate E:I ratio model was significantly associated with transmodality using an established measure of macroscale cortical organization(*42*). **b)** Nodal weights were also specifically correlated with the spatial patterns of benzodiazepine (BZD) sensitive GABA_A_ receptor subunit expression. Spatial relationships were tested for significance against a spatial-autocorrelation-preserving null distribution (BrainSMASH (*43)*) and corrected for multiple comparison using the Bonferroni correction (*p_Bonf_*).

### Development of the E:I ratio during adolescence

We next utilized our empirically generated E:I ratio model to test the hypothesis that E:I ratio declines as part of the critical period of association cortex development. An independent sample of 879 youth (aged 8-21.7 years) participated in a highly similar fMRI acquisition on the same scanner; this data was preprocessed using an identical pipeline. We applied our validated E:I model to the developmental dataset without further tuning and obtained the model-estimated distance from the classification hyperplane. This metric reflects a participant’s position on the continuum between “drug-like” (lower E:I) and “placebo-like” (higher E:I). To capture both linear and nonlinear effects in a rigorous statistical framework, we then regressed this metric on age using a generalized additive model with penalized splines (**Figure 1**, purple pathway). We found that age was positively associated with patterns of GABA-modulated functional connectivity, reflecting an age-related reduction in E:I ratio. Significant reductions occurred between ages 12.9 and 16.7 years (*F_s (Age)_* = 3.11, *p* = .037; **Supplemental Table 1**, “All connections”). The age-related reduction in E:I ratio was robust across multiple alternative parcellation schemes (**Supplemental Table 2**; **Supplemental Figure 2**).

### Age-related reductions in the E:I ratio are specific to association cortex

We hypothesized that age-related reductions in E:I ratio during adolescence were specific to association cortices. To test this hypothesis, we trained two additional models that restricted input features to connections to the most transmodal (**Figure 4a**, blue) or unimodal (**Figure 4a**, green) parts of the cortex. Both models significantly distinguished drug from placebo phMRI sessions (**Figure 4a**). However, when applied to the developmental dataset, significant age-related reductions in the E:I ratio were only observed for the model trained on connections with transmodal cortex (transmodal: *F_s (Age)_* = 9.96, *p* = .0017; unimodal: *F_s (Age)_* = 3.59, *p* = .058; transmodal vs. unimodal: *F_s (Age)_* = 5.96, *p* = .015; **Figure 4b**). These results suggest that transmodal association cortices undergo a reduction in E:I ratio during adolescence, consistent with a critical period of development.

**Figure 4.**
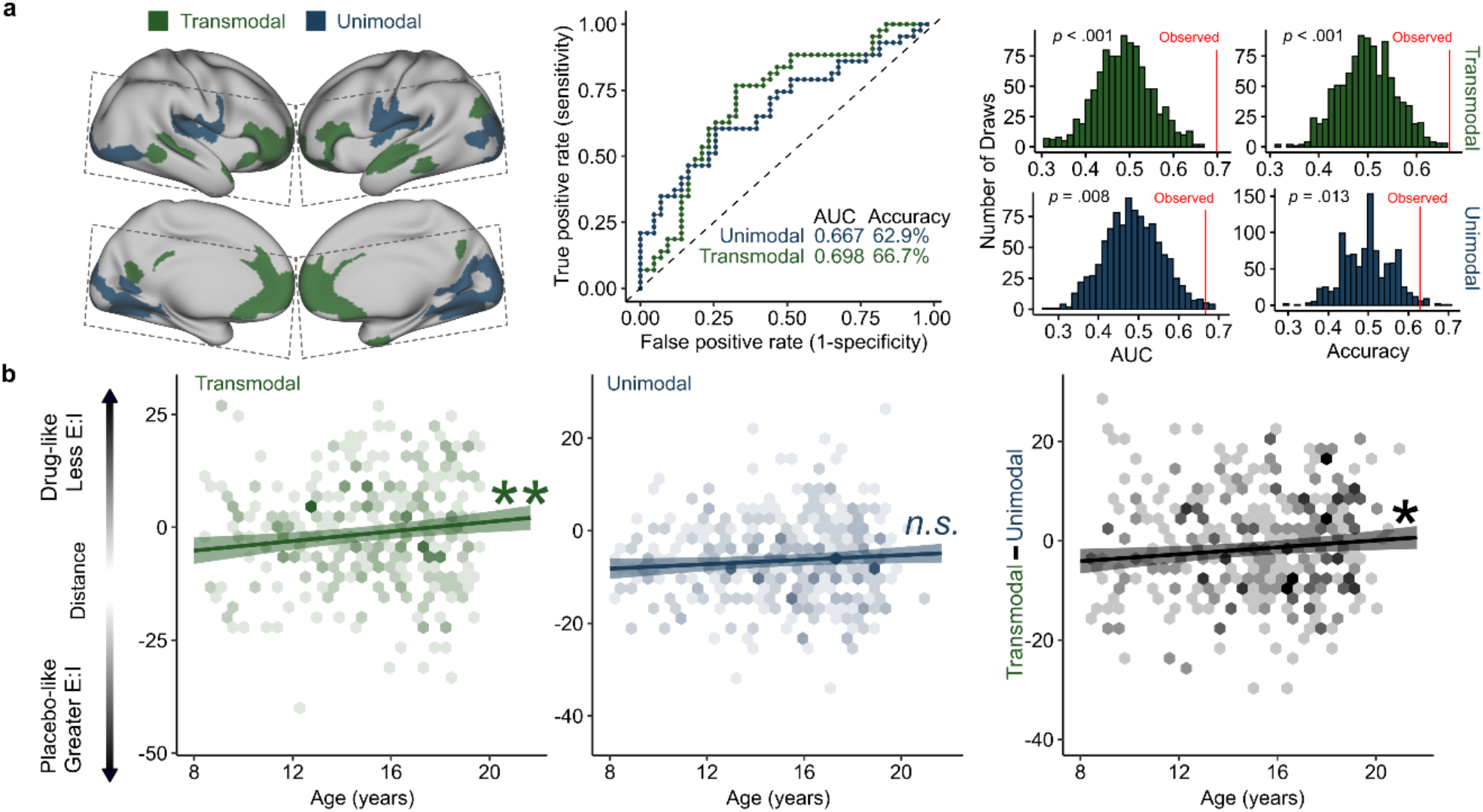
Transmodal areas undergo E:I ratio development during adolescence. **a)** Model performance for unimodal and transmodal classifiers. SVM classifiers were trained and validated for connections to the most transmodal (green) and most unimodal (blue) areas only. Dashed lines indicate acquisition field of view for the phMRI dataset. Both models performed significantly better than a permuted null distribution (middle: ROC curves for each model; right: null distributions from 1,000 null permutations). **b)** Models trained on transmodal and unimodal data were applied to the developmental dataset, generating a distance metric for each participant where greater values represent patterns of functional connectivity consistent with a lower E:I ratio. Individuals had lower estimated E:I ratio with age in transmodal cortex (left) but not in unimodal cortex (center). This pattern was confirmed by a significant effect of age on within subject change in transmodal vs. unimodal distance scores (right). \**p*<.05, \*\**p*<.01, *n.s*. not significant.

### Analysis of dimensions of psychopathology

Finally, we investigated whether individual differences in dimensions of psychopathology (*49*) were associated with the E:I ratio of association cortex. We found that mood disorder symptomatology, but not other psychopathology dimensions, moderated age-related differences in estimated transmodal E:I ratio. Specifically, individuals with greater lifetime mood disorder symptoms displayed a relatively stable E:I ratio over development instead of the normative reduction of the E:I ratio (Age*Mood interaction: *F* = 7.64, *p* = .0058).

## Discussion

We utilized a unique combination of human phMRI and developmental fMRI data to provide novel evidence for an essential component of critical period development: developmental reductions in the E:I ratio. Our approach generated an empirical fMRI model of the E:I ratio that showed a high degree of correspondence to known GABAergic benzodiazepine neuropharmacology and which could be applied to a large, independent sample of youth. Consistent with our hypothesis, this approach revealed that patterns of functional neurodevelopment in adolescence are consistent with developmental reductions in the E:I ratio that are specific to association cortex. Further, we show that individual differences in this process are associated with individual differences in lifetime mood symptom burden, in alignment with models positing that E:I abnormalities underlie the emergence of psychopathology (*9*, *29*, *30*, *50*–*52*). Together, these findings support the hypothesis that critical period mechanisms shape association cortices during adolescence.

Critical period development has been predominantly associated with early sensory cortex development. Since the first studies of critical period development in the visual cortex almost 60 years ago (*53*), a wealth of prior work has elucidated the mechanisms that shape critical period plasticity in these areas. These studies have identified the maturation of local inhibitory circuitry, particularly PV interneurons, and its resulting impact on the E:I balance as an essential critical period mechanism (*13*, *16*). This phenomenon is necessary for the opening of the critical period window, facilitates critical period plasticity, and is present in critical periods across sensory modalities (*16*, *17*, *23*, *37*). For example, modulation if the E:I balance using benzodiazepines has been shown to be sufficient to accelerate the timing of critical period plasticity (*13*, *37*). The results of this study suggest that this phenomenon also occurs in association cortex during human adolescence.

Our findings align with a growing literature characterizing inhibitory maturation during this developmental stage. Animal models and post mortem human studies have shown maturation of inhibitory neurobiology in the prefrontal cortex during adolescence, including increasing expression of PV interneurons and GABA_A_ α1 receptor subunits (*19*, *20*, *24*, *25*). These processes increase functional inhibition, reducing the E:I ratio and increasing the signal-to-noise ratio of circuit activity (*16*, *27*, *54*). Computational simulations have suggested that these maturational also facilitate high-frequency oscillatory capability (*27*). This is consistent with human EEG studies showing increased gamma-band oscillatory power during adolescence (*55*, *56*). Finally, two recent magnetic resonance spectroscopy studies have showed increases in GABA levels relative to glutamate levels in frontal cortex during adolescence (*57*, *58*). Though it is not possible to examine the functional effect of these changes on the E:I ratio using spectroscopy, these findings align with a model of developmental reduction in the E:I ratio during adolescence. This body of prior work cohere with the findings presented here, and are consistent with a critical period model of adolescent association cortex development. Just as sensory critical period plasticity refines neural circuits underlying sensory processing, the critical period for association cortex may facilitate the plasticity of circuits that underlie the higher-order cognitive processes refined during adolescence and are thought to be dependent on association cortex (*2*, *9*, *59*).

It should be noted that there are two classes of critical period mechanisms: *Facilitating factors* which open the critical period window and facilitate plasticity, and *braking factors* which stabilize neural circuits and physically limit future plasticity (*11*, *13*). The maturation of inhibition and the resulting reduction in the E:I ratio are critical period facilitators(*11*). Using generalized additive models, which can flexibly capture linear and nonlinear effects while penalizing overfitting, we found that the model fit for age-related reductions in the association cortex E:I ratio was linear. It is important to note that this does not necessarily mean that critical period plasticity is linearly increasing or that the critical period window is persistently open over the entire age range reported here. The developmental reduction in the E:I ratio is indicative of critical period opening, but it does not provide information about critical period closure. Closure of the adolescent critical period would be dependent upon the development of braking factors, such as myelination and the formation of perineuronal nets (PNN)(*60*–*62*), which may follow distinct developmental trajectories. Consistent with a critical period model, many studies have provided evidence of myelination of association cortex and large white matter pathways linking association cortex to other areas of the brain that continues into adulthood, including histological (*63*), myelin mapping (*64*–*66*), and diffusion imaging (*67*, *68*). At present, studies of PNN formation are limited to postmortem methods and animal models which have demonstrated developmental increases in PNN formation in the prefrontal cortex from adolescence to adulthood (*69*, *70*). Together, these studies indicate that critical period braking factors are forming during the transition from adolescence to adulthood, stabilizing neural circuits and closing the critical period window. However, in order to precisely demarcate the opening and closing of critical period plasticity during adolescence, future work is needed that jointly investigates the developmental timecourse of critical period facilitators, such as the E:I ratio reported here, and critical period braking factors.

Mood-related psychopathology typically first emerges during adolescence, with adolescent onset predicting greater illness chronicity and comorbidity (*71*, *72*). Here, we observed that beginning in adolescence, youth with greater burden of mood symptoms exhibit an altered trajectory of E:I development within the association cortex. Specifically, greater lifetime mood symptom burden was associated with reduced development of inhibition in transmodal regions of the brain. Many studies have linked the occurrence of psychopathology with E:I disruption (*29*, *30*, *50*, *73*, *74*), and cross-species research has specifically implicated GABA-mediated E:I disruptions in the etiology of mood psychopathology (*75*, *76*). Specifically, animal studies have shown that initial reductions in GABAergic inhibition lead to downstream reductions in glutamatergic transmission, and to alteration of the normal E:I balance (*30*, *50*). Human studies have provided convergent evidence, demonstrating reduced GABA levels in the brain in those with depression(*77*)_as well as reduced glutamate in individuals with more severe anhedonia (*78*). Conversely, the pharmacologic enhancement of GABAergic signaling within association regions has been shown to have antidepressant effects (*30*). As such, our study supports the hypothesis that the pathophysiology of depression in part involves altered glutamatergic and GABAergic signaling (*30*, *50*). Moreover, it places this hypothesis within a neurodevelopmental framework—underscoring how E:I disruptions can manifest due to atypical critical period development.

Finally, we note that the approach used in this study highlights the potential for phMRI data to generate insights into independent datasets to inform new hypotheses. We combined machine learning and phMRI using a GABAergic alprazolam challenge to generate an empirical model for the effect of GABAergic modulation on patterns of fMRI connectivity. As evidence for the efficacy of this approach, the trained model could not only significantly predict drug versus placebo sessions in unseen data, but the model features demonstrated a significant correspondence with known benzodiazepine neuropharmacology. Notably, the model features were significantly associated with the GABA receptor most strongly implicated in critical period development, the GABA_A_ α1 receptor. Though, due to the rarity of phMRI data, we were not able to confirm the generalizability of our model with an independent alprazolam phMRI dataset, the model performance and underlying interpretability of the learned features highlights the biological relevance of this method. Whereas in this study we applied this method to an independent developmental dataset to provide insights into the critical period mechanisms unfolding during adolescence, future work could apply this approach to other datasets to inform new research questions.

Together, these findings support the hypothesis that critical period mechanisms, such as the inhibition-induced reduction of the E:I ratio, shape association cortices during adolescence. Studying development from a critical period perspective provides a powerful mechanistic framework for understanding how experience and neurobiology interact to shape long-term cognitive, social, and psychiatric outcomes. Importantly, a critical period model of adolescent development can draw on the history of detailed work on sensory critical periods to generate testable hypotheses for the mechanisms unfolding during adolescence in association cortex. Understanding these mechanisms are a necessary prerequisite to understanding of how experience, environment, and neurobiology contribute to differing neurodevelopmental trajectories in health and mental illness. This work thus lays the groundwork for future studies of the unique impact of experience on neurodevelopment and also suggests the possibility of targeted interventions during this critical window of vulnerability to psychopathology (*79*).

## Materials and Methods

### Participants and experimental procedures

#### Alprazolam sample

The alprazolam sample and study procedures have been described in detail in our earlier work(*80*). Briefly, forty-seven adults participated in a double-blind, placebo controlled pharmacological imaging study using the benzodiazepine alprazolam. Each participant completed two identical experimental sessions approximately one week apart. In one session, participants were given a 1 mg dose of alprazolam, and in the other they were given an identical appearing placebo. The order of administration was counter-balanced across participants. During both sessions, participants completed an emotion identification task that lasted 10.5 minutes while functional MRI (fMRI) data was collected. Task-related fMRI results have been previously reported(*80*). Four participants were excluded due to excess head motion in at least one session (see below) for a final sample of 43 participants and 86 sessions total (ages 20.9 - 59.4; *M* = 40.3, *SD* = 13.12, male/female = 24/19). Study procedures were approved by the University of Pennsylvania IRB, and all participants provided written informed consent.

#### Developmental sample

Neuroimaging data were obtained from a community-based sample of 1,476 youth (ages 8 – 21.9, *M* = 14.63, *SD* = 3.43; male/female = 698/778) that were part of the Philadelphia Neurodevelopmental Cohort (PNC). Data collection procedures and sample characteristics have been previously described in detail(*49*, *81*, *82*). Functional MRI data were collected while participants performed the same emotion identification task as the alprazolam sample; this is also described in previous work(*81*). From this original sample, 306 participants were excluded based on health criteria, including psychoactive medication use at the time of study, medical problems that could impact brain function, a history of psychiatric hospitalization, and gross structural brain abnormalities. A total of 234 participants were excluded from further analysis due to head motion (see below) and 56 were excluded for poor structural image quality. In sum, following health exclusions and rigorous quality assurance, we retained 879 participants (ages 8.0 - 21.7 at first visit, *M* = 14.95, *SD* = 3.24; male/female = 383/496).

### Neuroimaging acquisition

#### Alprazolam sample

All data were collected on a Siemens Trio 3T as previously reported(*80*). Whole-brain structural data were obtained with a 5-minute magnetization-prepared, rapid acquisition gradient-echo T1-weighted image (MPRAGE) using the following parameters: TR 1620ms, TE 3.87 ms, field of view (FOV) 180×240 mm, matrix 192×256, effective voxel resolution of 1 x 1 x 1mm. BOLD fMRI data were obtained as a slab single-shot gradient-echo (GE) echoplanar imaging (EPI) sequence using the following parameters: TR = 3000, TE = 32 ms, flip angle = 90°, FOV = 240 mm, matrix = 128 X 128, slice thickness/gap = 2/0mm, 30 slices, effective voxel resolution of 1.875 x 1.875 x 2mm, 210 volumes. As previously described(*80*), data were acquired in a FOV that included temporal, inferior frontal, and visual cortices as well as subcortical structures (Figure 3a; gray boxes).

#### Developmental sample

All neuroimaging data were collected on the same Siemens Trio 3T scanner as was used for the alprazolam dataset. The neuroimaging procedures and acquisitions parameters have been previously described in detail(*81*). Briefly, structural MRI was acquired with a 5-min MPRAGE T1-weighted image (TR = 1810 ms; TE = 3.51 ms; TI = 1100 ms, FOV = 180 × 240 mm^2^, matrix = 192 × 256, effective voxel resolution = 0.9 × 0.9 × 1 mm^3^). BOLD fMRI was acquired using similar acquisition parameters to the alprazolam dataset. BOLD fMRI scans were acquired as single-shot, interleaved multi-slice, GE-EPI sequence sensitive to BOLD contrast with the following parameters: TR = 3000 ms, TE = 32 ms, flip angle = 90°, FOV = 192 × 192 mm^2^ (whole brain acquisition), matrix = 64 × 64; 46 slices, slice thickness/gap = 3/0 mm, effective voxel resolution = 3.0 × 3.0 × 3.0 mm^3^, 210 volumes.

### Preprocessing of neuroimaging data

All preprocessing was performed using fMRIPrep 20.0.7 (RRID:SCR_016216;(*83*), which is based on Nipype 1.4.2(*84*), and XCP Engine (PennBBL/xcpEngine: atlas in MNI2009 Version 1.2.3; Zenodo: http://doi.org/10.5281/zenodo.4010846; (. The neuroimaging data from the alprazolam and developmental datasets were processed using identical pipelines as described below.

#### Anatomical data preprocessing

The T1-weighted (T1w) image was corrected for intensity non-uniformity (INU) with N4BiasFieldCorrection(*85*), distributed with ANTs 2.2.0(*86*), and used as T1w-reference throughout the workflow. The T1w-reference was then skull-stripped with a Nipype implementation of the antsBrainExtraction.sh workflow (from ANTs), using OASIS30ANTs as target template. Brain tissue segmentation of cerebrospinal fluid (CSF), white-matter (WM) and gray-matter (GM) was performed on the brain-extracted T1w using FAST in FSL 5.0.9(*87*). Volume-based spatial normalization to MNI2009c standard space was performed through nonlinear registration with antsRegistration (ANTs 2.2.0), using brain-extracted versions of both the T1w reference and the T1w template.

#### Functional data preprocessing

The alprazolam dataset consisted of two BOLD acquisitions per participant (drug and placebo session) which were preprocessed individually. The developmental dataset consisted of one BOLD acquisition per participant. All BOLD acquisitions were processed with the following steps. BOLD runs were first slice-time corrected using 3dTshift from AFNI 20160207(*88*) and then motion corrected using mcflirt (FSL 5.0.9;(*87*). A fieldmap was estimated based on a phase-difference map calculated with a dual-echo GRE sequence, processed with a custom workflow of SDCFlows inspired by the <u>epidewarp.fsl script</u> and further improvements in HCP Pipelines(*89*). The fieldmap was then co-registered to the target EPI reference run and converted to a displacement field map with FSL’s fugue and other SDCflows tools. Based on the estimated susceptibility distortion, a corrected BOLD reference was calculated for a more accurate co-registration with the anatomical reference. The BOLD reference was then co-registered to the T1w reference using bbregister (FreeSurfer) which implements boundary-based registration(*90*). Co-registration was configured with nine degrees of freedom to account for distortions remaining in the BOLD reference. Six head-motion parameters (corresponding rotation and translation parameters) were estimated before any spatiotemporal filtering using mcflirt. Finally, the motion correcting transformations, field distortion correcting warp, BOLD-to-T1w transformation and T1w-to-template (MNI) warp were concatenated and applied to the BOLD timeseries in a single step using antsApplyTransforms (ANTs) with Lanczos interpolation.

Confounding time-series were calculated based on the preprocessed BOLD data. The global signal was extracted within the whole-brain mask. Additionally, a set of physiological regressors were extracted to allow for component-based noise correction (CompCor, Behzadi et al. 2007). Anatomical CompCor (aCompCor) principal components were estimated after high-pass filtering the preprocessed BOLD time-series (using a discrete cosine filter with 128s cut-off). The aCompCor components were calculated within the intersection of the aforementioned mask and the union of CSF and WM masks calculated in T1w space, after their projection to the native space of each functional run (using the inverse BOLD-to-T1w transformation). Components were also calculated separately within the WM and CSF masks. In this study, for each aCompCor decomposition, the k components with the largest singular values were retained, such that the retained components’ time series were sufficient to explain 50 percent of variance across the nuisance mask (CSF and WM). The remaining components were dropped from consideration. The head-motion estimates calculated in the correction step were also placed within the corresponding confounds file. The confound time series derived from head motion estimates and global signals were expanded with the inclusion of temporal derivatives and quadratic terms for each(*91*).

Subject-level timeseries analysis was carried out in XCP Engine using FILM (FMRIB’s Improved General Linear Model)(*92*). All event conditions from the emotion identification task(*80*, *81*) were modeled in the GLM as 5.5 second boxcars convolved with a canonical hemodynamic response function. Each of the five emotions (fear, sad, angry, happy, neutral) was modeled as a separate regressor. The temporal derivatives and quadratic terms for each task condition as well as the confounding aCompCor, global signal, and motion timeseries described above were included as nuisance regressors. Task regression has been shown to produce patterns of BOLD fMRI connectivity that are highly similar to those present at rest(*93*), and convergent results from several independent studies that have shown that functional networks are primarily defined by individual-specific rather than task-specific factors (Gratton et al., 2018). The nuisance regression pipeline used here has been shown be a top-performing procedure for mitigating motion artifacts(*41*). Consistent with our prior work, participants in the alprazolam dataset were excluded from future analyses if mean framewise displacement exceeded 0.5 mm in either session. A more stringent threshold of 0.3 mm was applied to the developmental dataset; head motion was also included as a covariate in all developmental models (see below).

### Connectivity matrix generation

Fully preprocessed fMRI data were used to generate mean timeseries within a set of atlas-defined brain regions for each participant. Cortical regions were defined according to the Schaefer 400 parcel cortical atlas(*94*). To accommodate the restricted FOV of the alprazolam BOLD acquisition, the atlas was masked such that only parcels with greater than 95% coverage were included in connectivity analyses. Subcortical regions were defined using the Automated Anatomical Labeling (AAL) atlas(*95*). Subcortical areas included the left and right caudate, putamen, accumbens, pallidum, thalamus, amygdala, hippocampus, and parahippocampal area. These cortical and subcortical atlases were combined and used to generate mean timeseries for each region in each dataset. Functional connectivity was calculated as the correlation coefficient of the timeseries for each pair of regions (20,503 unique pairs). As part of sensitivity analyses, we repeated this process after defining cortical areas using Schaefer 200 parcellation(*94*), the Multi-modal Parcellation atlas(*96*), the Gordon cortical atlas(*97*), or the AAL(*95*).

### Pharmacological classification analysis

We used a linear support vector machine (SVM) to classify drug vs. placebo sessions in the alprazolam dataset based on multivariate patterns of functional connectivity. Linear SVMs find a hyperplane to separate two classes of data by maximizing the margin between the closest points (the support vectors; (. SVMs were implemented in R using the e1071 library(*99*) and were trained using a linear kernel and the default parameters. Model performance was evaluated using 10-fold cross-validation, iteratively selecting data from 90% of participants as training data and testing the trained model on data from the remaining 10% of participants. Across testing sets, the prediction accuracy and area under the receiver operating curve (AUC) were calculated to evaluate model performance. To ensure our results were not driven by a specific cross-validation split, we repeated the entire 10-fold cross-validation procedure 100 times, drawing the 10-fold subsets at random each time. Performance metrics were finally averaged across the 100 iterations of the cross-validation procedure.

To evaluate if model performance (i.e., the accuracy and the AUC) was significantly better than expected by chance, we performed a permutation test(*100*). Specifically, we re-applied the cross-validation procedure 1,000 times, each time permuting the session labels (drug and placebo) across the training samples without replacement. Significance was determined by ranking the actual prediction accuracy versus the permuted distribution; the *p*-value of the accuracy and AUC was calculated as the proportion of permutations that showed a higher value than the observed value in the real, unpermuted data.

#### Analysis of feature weights

After cross-validation and significance testing, we trained the model on all participants and extracted the feature weights for further analysis. First, we calculated the absolute value of the weights and summed them across all connections (edges) for a given region (node) to compare the overall contribution of each region to the model, irrespective of the sign of the feature weights(*100*). Next, to evaluate the spatial pattern of the feature weights, we calculated the mean signed feature weight for each node, reflecting the directionality of the effect of the drug manipulation according to the trained model. We then used this feature map to assess the biological relevance of our trained model. Specifically, we calculated the spatial correlation between this pattern of nodal feature weights with two sets of cortical features. The first was the widely used principal gradient of macroscale cortical organization(*42*), which places each cortical region on a continuum between unimodal (i.e. sensorimotor cortices) to transmodal (i.e. association cortices) function. The second set of cortical features was selected based on the known pharmacology of benzodiazepines like alprazolam. Alprazolam is a positive allosteric modulator of the GABA_A_ receptor, and of the six GABA_A_ α subunits (α1-6), only subunits α1, α2, α3, and α5 are benzodiazepine sensitive (*48*, *101*). To quantify the spatial distribution of the six GABA_A_ α subunits, we extracted the microarray gene expression patterns for their corresponding GABA_A_ receptor genes (GABR_A_1-6) from the Allen Human Brain Atlas (data available at https://www.meduniwien.ac.at/neuroimaging/mRNA.html) (*46*, *102*). For each of the six gene expression maps, we quantified the mean expression value within each cortical parcel and calculated the spatial correlation with the pattern of nodal SVM weights.

To test the significance of the spatial correlation between our pattern of cortical feature weights and each of the biological brain maps, we compared the observed correlation value to a null distribution generated with BrainSMASH (Brain Surrogate Maps with Autocorrelated Spatial Heterogeneity; https://brainsmash.readthedocs.io/; (. The spatial autocorrelation of brain maps can lead to inflated *p*-values in spatial correlation analyses and must be accounted for in the creation of null models. BrainSMASH addresses this by generating permuted null brain maps that match the spatial autocorrelation properties of the input data. We used BrainSMASH to generate 10,000 spatial-autocorrelation-preserving null permutations based on the input data and the pairwise distance matrix for the cortical parcellation, generating a null distribution of spatial correlation coefficients. We calculated two-tailed *p*-values by squaring all correlation values (i.e. spatial *R*^2^) and calculating the proportion of times the null distribution exceeded the observed value.

#### Transmodal and unimodal classification models

Our primary hypothesis was that E:I ratio reductions would be specific to association cortices during youth. In order to test this hypothesis directly, we trained two additional models after applying an *a priori* feature selection step. Specifically, we thresholded the top and bottom quartiles of cortical parcels based on their position in the principal gradient of functional organization (*42*), with the top 25% representing transmodal association cortex and the bottom 25% representing unimodal sensory cortex. We then created two new feature sets that restricted the input features to connections to these transmodal or unimodal areas only. This selection procedure ensured that the resulting numbers of features were equal between the two feature sets (9,541 features per model). We then trained and validated the transmodal and unimodal models according the procedures described above.

### Developmental analyses

#### Application of the pharmacological model to the developmental dataset

After training and validating the pharmacological benzodiazepine models, we applied the models to the functional connectivity data for each participant in the developmental sample. For each participant, each model yielded the distance from the classification hyperplane that separates the two classes (drug vs. placebo). Observations close to the hyperplane (distance values near zero) are less representative of the class, and those further from the hyperplane are more representative. The distance metric is such that values greater than zero indicate more “drug-like” patterns of functional connectivity and values less than zero indicate more “placebo-like” patterns of connectivity. As the pharmacological effect of alprazolam is to increase GABAergic inhibitory signaling, more “drug-like” patterns reflect greater GABA-ergic inhibitory modulation of functional connectivity. As such, more “drug-like” patterns were interpreted to reflect a reduced E:I balance relative to more “placebo-like” patterns. These distance metrics were normally distributed and thus provided a continuous measure of E:I balance for use in further analyses. We first applied the model trained on all the input features and then applied the transmodal- and unimodal-specific models, generating three sets of distance values per participant.

#### Developmental regression models

To assess the developmental trajectory of E:I balance, we modeled the classification distance metrics from each model as a function of age using penalized splines within a generalized additive model (GAM). GAMs allow us to flexibly capture linear or nonlinear age effects while penalizing overfitting. To test for windows of significant change across the age range, we calculated the first derivative of the smooth function of age from the GAM model using finite differences and then generated a simultaneous 95% confidence interval of the derivative following the method described by Simpson (and implemented using the *gratia* library(*103*) in R. Intervals of significant change were identified as areas where the simultaneous confidence interval of the derivative does not include zero. To test if the effect of age on classification distance differed between the transmodal and unimodal SVM models, we calculated the residualized change (*104*) in transmodal vs. unimodal distance scores by regressing the unimodal distance out of transmodal distance. We then regressed the residualized change score on age using a GAM. All models included sex as a covariate as well as head motion and attentiveness as covariates of no interest. Head motion was quantified as mean framewise displacement during the fMRI acquisition. Attentiveness was quantified as the number of response omissions during the emotion identification task; this covariate was included to control for potential effects of arousal on model performance as alprazolam can cause drowsiness. All GAMs were fit using the mgcv library(*105*) in R.

### Analysis of dimensions of psychopathology

As previously described (*49*, *106*, *107*), PNC participants underwent a clinical assessment of psychopathology. Multiple domains of psychopathology symptoms were evaluated using a structured screening interview (GOASSESS); we used this data to investigate whether dimensions of psychopathology moderated developmental reductions in E:I balance. As has been detailed in prior work (*49*, *106*, *107*), factor scores were derived from the clinical assessments using a bifactor confirmatory factor analysis model that included a general factor for overall psychopathology as well as four specific factors that primarily represent anxious-misery (mood & anxiety) symptoms, psychosis-spectrum symptoms, behavioral symptoms (conduct and ADHD), and fear symptoms (phobias). Importantly, all five factors are orthogonal and can be considered jointly in analysis of imaging data. In order to sample a broad range of factor scores, we expanded our inclusion criteria to include individuals with a history of psychiatric hospitalization (*N* = 1018; ages 8 – 21.7; *M* = 15.0, *SD* = 3.23, male/female = 462/556). We analyzed these data in a GAM that included age-by-factor score interactions for each factor from the bifactor model. Interactions were fit as bivariate smooth interactions with penalized splines using tensor interaction smooths (‘ti’ in mgcv).

### Data and code availability

The developmental dataset is publicly available in the Database of Genotypes and Phenotypes (dbGaP accession <u>phs000607.v3.p2</u>). Pharmacological imaging data is available upon reasonable request. All code used for pharmacological classification analyses and developmental analyses are available at https://pennlinc.github.io/Larsen_EI_Development/.

## Author contributions

B.L. and T.D.S. conceived and designed the analyses. B.L., Z.C., and A.A. analyzed the data. J.S., A.P., M.B., and T.M.M contributed analysis tools. A.A.B., D.R.R., V.J.S., R.E.G., R.C.G., A.S.M., M.E.C., A.P., and M.B. commented on analyses. B.L. wrote the paper with contributions from all authors. T.D.S. and D.H.W. supervised the work.

## Supplemental Data

**Supplemental Table 1.**
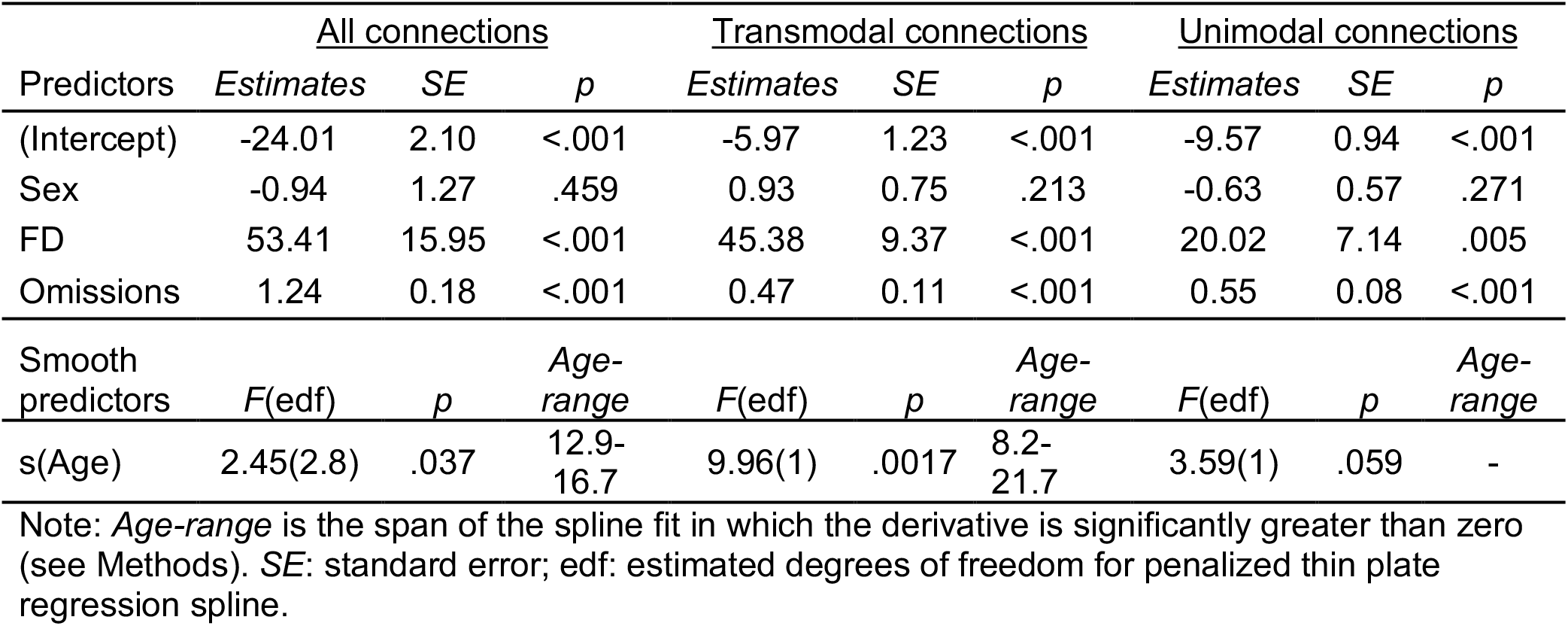
Regression table for developmental generalized additive models (GAM).

**Supplemental Table 2.**
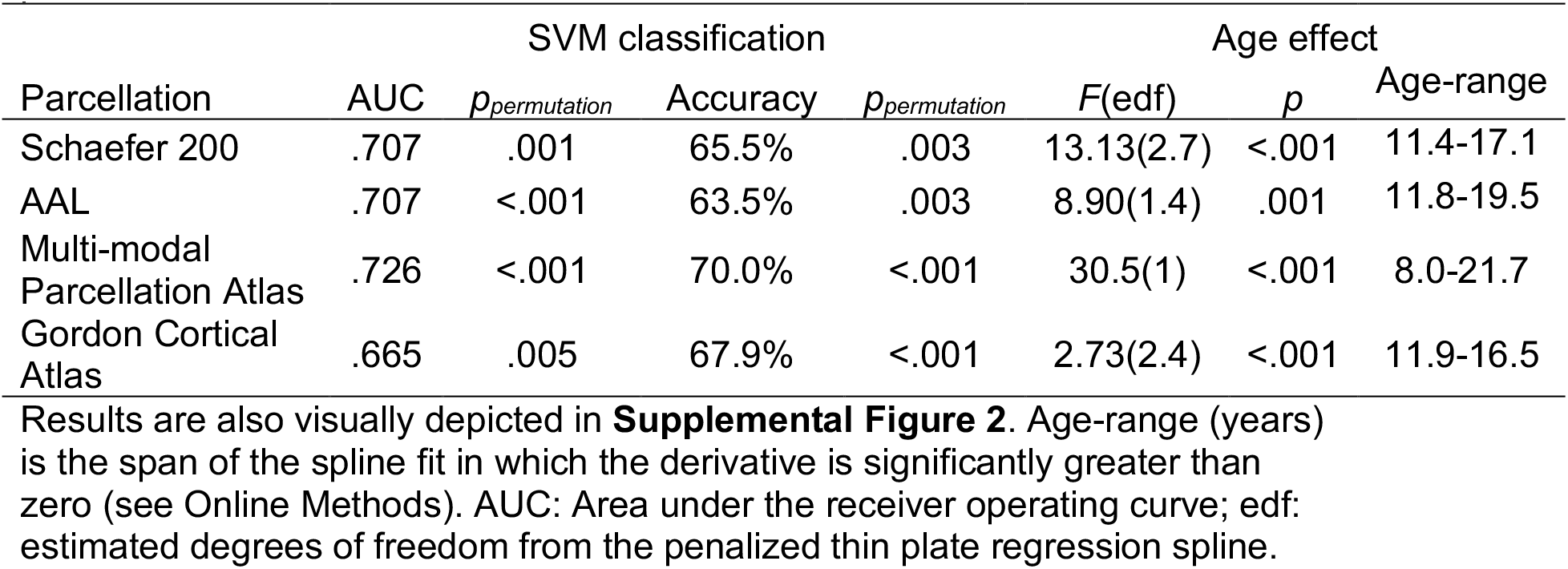
Classification results and age effects for alternative parcellation atlases.

**Supplemental Figure 1.**
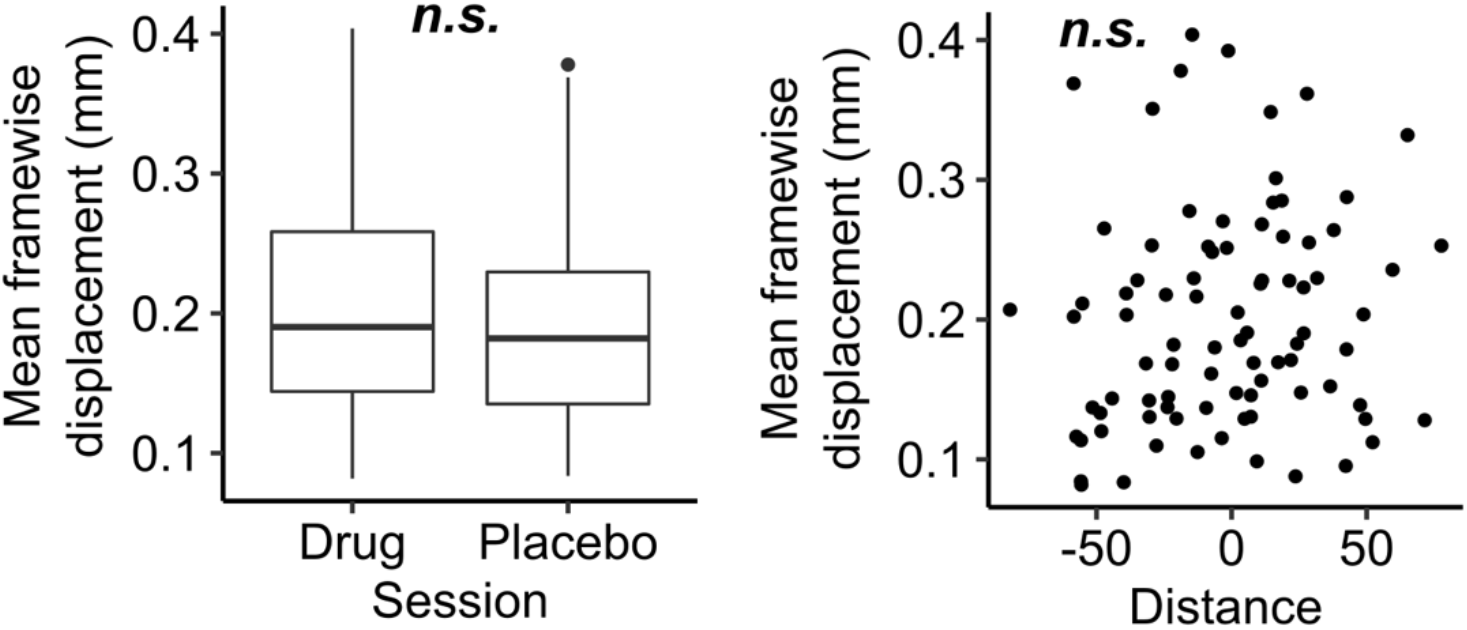
Head motion is not associated with the pharmacological manipulation or classifier performance. Left: A summary of in-scanner head motion (mean framewise displacement) did not differ between drug and placebo sessions (*t_paired_* = 1.44, *p* = .16). Right: The same metric of head motion was also not associated with model-predicted distance from the classification plane (*r* = .13, *p* = .24).

**Supplemental Figure 2.**
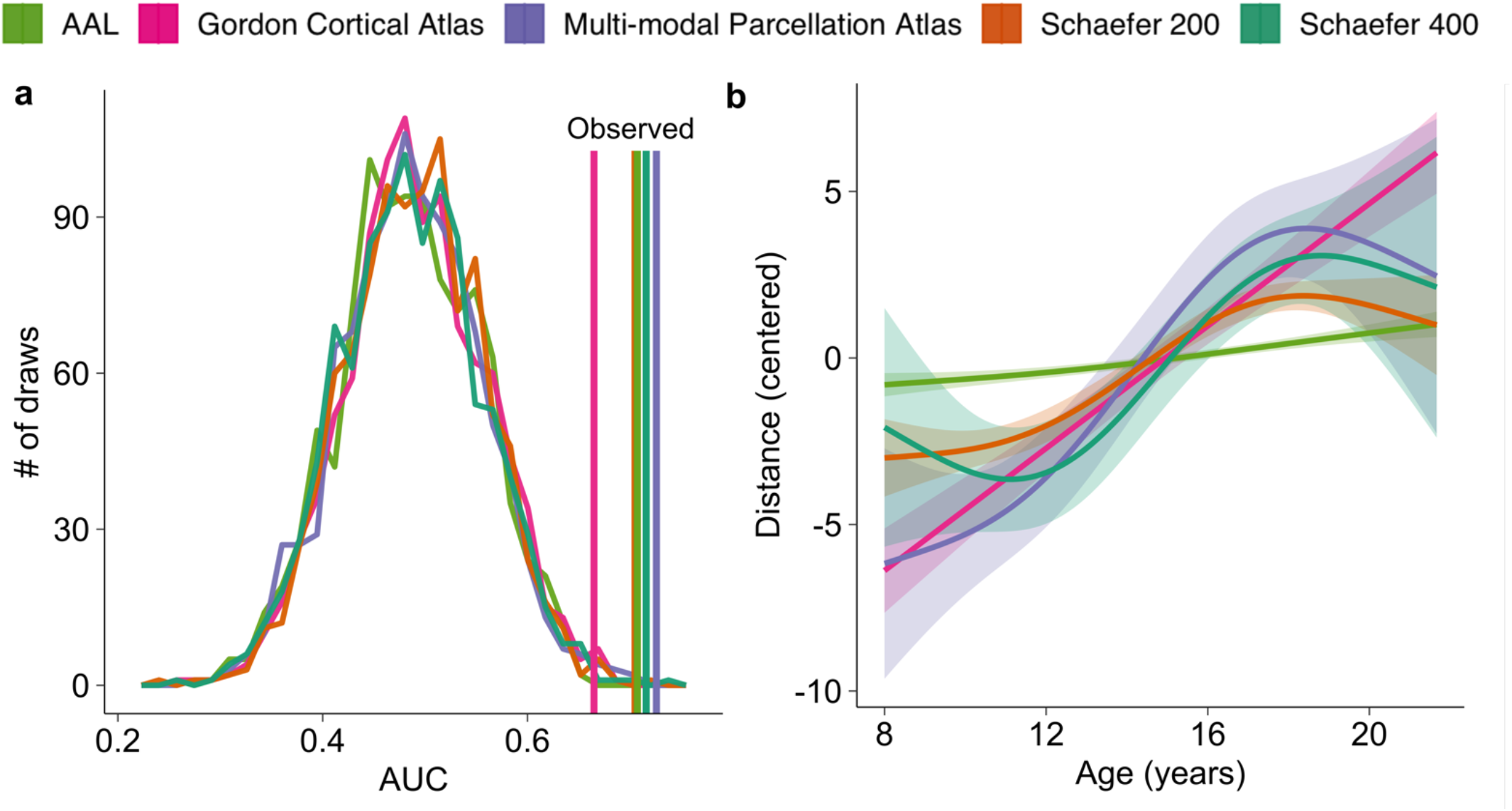
Classification and developmental effects are robust to alternative cortical parcellation schemes. **a)** Classifier performance. All binary SVM classifiers identified drug and placebo sessions in 10-fold cross-validation with an AUC values that exceeded the randomly permuted null AUC distributions (see **Supplemental Table 1)**. **b)** When the models from panel **a)** were applied to the developmental dataset, models trained on all parcellation schemes produced distance metrics that significantly increased with age during adolescence, indicating age-related reductions in the E:I ratio. Distance metrics are centered to facilitate comparisons between the model predictions. GAM model statistics are reported in **Supplemental Table 1**.

